# Collaborative role of two distinct cilium-specific cytoskeletal systems in driving Hedgehog-responsive transcription factor trafficking

**DOI:** 10.1101/2024.09.26.615198

**Authors:** Pei-I Ku, Jamuna S Sreeja, Abhishek Chadha, David S Williams, Martin F Engelke, Radhika Subramanian

**Author notes:** These authors contributed equally: Pei-I Ku and Jamuna S Sreeja.

## Abstract

Calibrated transcriptional outputs in cellular signaling require fine regulation of transcription factor activity. In vertebrate Hedgehog (Hh) signaling, the precise output of the final effectors, the GLI (Glioma-associated-oncogene) transcription factors, depends on the primary cilium. In particular, the formation of the activator form of GLI upon pathway initiation requires its concentration at the distal cilium tip. However, the mechanisms underlying this critical step in Hh signaling are unclear. We developed a real-time imaging assay to visualize GLI2, the primary GLI activator isoform, at single particle resolution within the cilium. We observed that GLI2 is a cargo of Intraflagellar Transport (IFT) machinery and is transported with anterograde bias during a restricted time window following pathway activation. Our findings position IFT as a crucial mediator of transcription factor transport within the cilium for vertebrate Hh signaling, in addition to IFT’s well-established role in ciliogenesis. Surprisingly, a conserved Hh pathway regulator, the atypical non-motile kinesin KIF7, is critical for the temporal control of GLI2 transport by IFT and the spatial control of GLI2 localization at the cilium tip. This discovery underscores the collaborative role of a motile and a non-motile cilium-specific cytoskeletal system in determining the transcriptional output during Hh signaling.

## INTRODUCTION

The primary cilium, a solitary microtubule-based organelle located on the cell surface, plays a central role in receiving and transmitting a multitude of extracellular signals in a tissue-specific manner^1–5^. Understanding how the cilium enables precisely calibrated transcriptional output in response to the extracellular morphogen is a fundamental question in signal transduction. In vertebrates, the Hedgehog (Hh) pathway exhibits a strict dependence on the primary cilium^6,7^. Additionally, several key pathway components reside in the cilium^8–11^. As a result, Hh signaling has emerged as an important system for deciphering the mechanisms by which the cilium serves as a subcellular compartment for signal transduction.

Glioma-associated oncogene (GLI) transcription factors are the main effectors of Hh pathway^12,13^. Among them, isoforms GLI2 and GLI3 function as the main activator (GLI-A) and proteolytically cleaved repressor (GLI-R) forms, respectively^14–16^. The precise ratio of GLI-A to GLI-R determines the final transcriptional output in response to varying levels of Hh morphogen input^17^. In vertebrate Hh signaling, the primary cilium is required to generate the correct ratios of GLI-A and GLI-R^8,18–20^. A key event following Hh pathway activation is the localization of GLI proteins to the cilium tip, known as the “cilium tip compartment”, prior to translocating to the nucleus^8,11,20–22^. Aberrant cilium localization of GLI proteins is associated with incorrect transcriptional output^23–28^. Despite the fundamental importance of this process in Hh signaling, the mechanism by which GLI proteins are enriched at this sub-ciliary compartment is unknown. This void exists because the dynamics of GLI and its associated regulators have not been directly visualized due to the technical challenges of imaging low copy-number non-membrane bound Hh regulators in the small ciliary volume of mammalian cells. Consequently, the contribution of the ciliary cytoskeletal proteins to GLI regulation and, therefore, the cilium-dependence of the transcriptional output of Hh signaling remains unclear.

The conserved Intraflagellar Transport (IFT) system, which is composed of over 20 protein subunits, mediates the transport of several ciliary cargos, including tubulin^29,30^, and is essential for cilia formation and maintenance^31–35^. The anterograde movement of IFT to the cilium tip is powered by the KIF3A/3B motor proteins, while Dynein-2 motors drive the retrograde movement to the cilium base^36–39^. However, the precise role of IFT as a transport system in Hh signaling is challenging to establish. Inhibiting anterograde IFT components or the kinesin motors results in defective cilium assembly and consequently impairs Hedgehog signaling^6^. Inhibiting retrograde ciliary trafficking using IFT122 mutants or a Dynein-2 mutant or inhibitor results in abnormal accumulation of GLI2 within the cilium^40,41^. These interventions also compromise the structural integrity of the axoneme^40^. Furthermore, other Hh pathway components, such as SMO and PATCHED, which show incorrect cilium localization upon Dynein-2 inhibition^19,42,43^, do not exhibit stereotypical IFT-like dynamics^44–46^. Therefore, it is challenging to determine whether the Hh pathway defects observed in IFT mutants are due to irregular protein transport, the disruption of cilium integrity, or both. However, other evidence suggests that the impacts of IFT mutations may not be solely attributed to defects in the cilium structure. First, deleting two IFT components, IFT25 and IFT27, does not visibly affect the cilium structure but results in the failure to enrich GLI2 at the cilium tip in response to Hh pathway activation^25,47^. At present, it is not well-established whether the Hh signaling defects observed in the absence of IFT25/27 arise from their role in IFT transport of GLI proteins or another ciliary function for these IFT subunits^48^. Second, GLI2 has been found to associate with KAP3A, an adapter for the anterograde ciliary motor Kinesin-2^49^. Whether the interaction between KAP3A and GLI2 is related to cilium-specific GLI transport during Hh signaling is unknown. Thus, whether GLI proteins are transported within the cilium as an IFT cargo or through other active or diffusion-based processes remains unknown.

The ciliary kinesin-4 protein KIF7 (*Drosophila melanogaster* Costal2) is a conserved protein component of the Hh pathway^27,28,50^. Genetic inactivations and mutations of KIF7 lead to congenital malformations caused by mis-regulated Hh signaling in mice and humans^23,27,51–53,50,28^. KIF7 binds to GLI and accumulates at the cilium tip in response to Hh pathway activation^28,54^. In the absence of KIF7, GLI proteins fail to properly concentrate at the cilium tip^55,56^. Conversely, in the absence of GLI2 and GLI3, KIF7 does not accumulate at the cilium tip^54^. Together, these findings suggest that KIF7 and GLI act synergistically to establish a “Hh signaling cilium tip compartment”^54^. However, since KIF7 is a non-motile kinesin, it is not considered the transport motor for GLI in the cilium^57,58^. Currently, the precise roles of KIF7 in regulating GLI transport and localization at the cilium tip are poorly understood.

Here, we developed a total internal reflection fluorescence microscopy (TIRFM)-based assay for imaging GLI2, the primary activator form of GLI, in the cilium^20,59^. Single particle imaging established GLI2 as an IFT cargo. Upon pathway activation, GLI2 is preferentially loaded on anterograde IFT trains in a temporally restricted manner. Consistently, inhibition of anterograde IFT using an inhibitable KIF3A-KIF3B system prevents GLI2 accumulation at the cilium tip after Hh activation. Interestingly, we found that KIF7 plays an important role in regulating GLI2 ciliary transport in three ways: KIF7 restricts GLI2 loading on IFT before pathway activation, prevents aberrant localization of GLI2 along the cilium shaft, and is important for concentrating GLI2 at the cilium tip. This finding underscores the synergy between two cilium-specific cytoskeleton systems, one motile (IFT) and one non-motile (KIF7), in establishing a GLI2-enriched cilium tip compartment to enable precise Hh-responsive transcriptional output.

## RESULTS

### Real-time visualization of GLI2 accumulation at the cilium tip

To precisely monitor the spatiotemporal patterns of GLI2 accumulation within individual cilia in live cells over several hours, we established a multi-wavelength TIRFM assay (Fig. 1A). We used an established Flp-In^TM^ system for stable expression of N-terminal Halo-tagged GLI2 in NIH3T3 cells^60^. The responsiveness of this cell line to Hh ligands was verified by measuring *Gli1 and Ptch1* mRNA expression in response to the Smoothened agonist SAG21k (Supplementary Fig. 1A). For live imaging, cells were cultured in a low serum medium for 24 hours to promote cilia assembly (see Methods). Halo-tagged GLI2 was labeled with the JFX549 fluorophore^61^ following 24 hours of serum starvation to promote ciliogenesis and stimulate cilia growth. The cilium was visualized using stable expression of a ciliary membrane protein SSTR3-mNeonGreen (Fig. 1A, B). For multi-hour TIRFM imaging, we selected cilia that were oriented parallel to the coverslip and displayed limited mobility for 5-15 minutes before SAG21k activation. Imaging was performed at 1–2-minute intervals after adding SAG21k (as shown in Fig. 1B). This method allowed us to track GLI2 accumulation in individual cilia over many hours.

**Figure 1.**
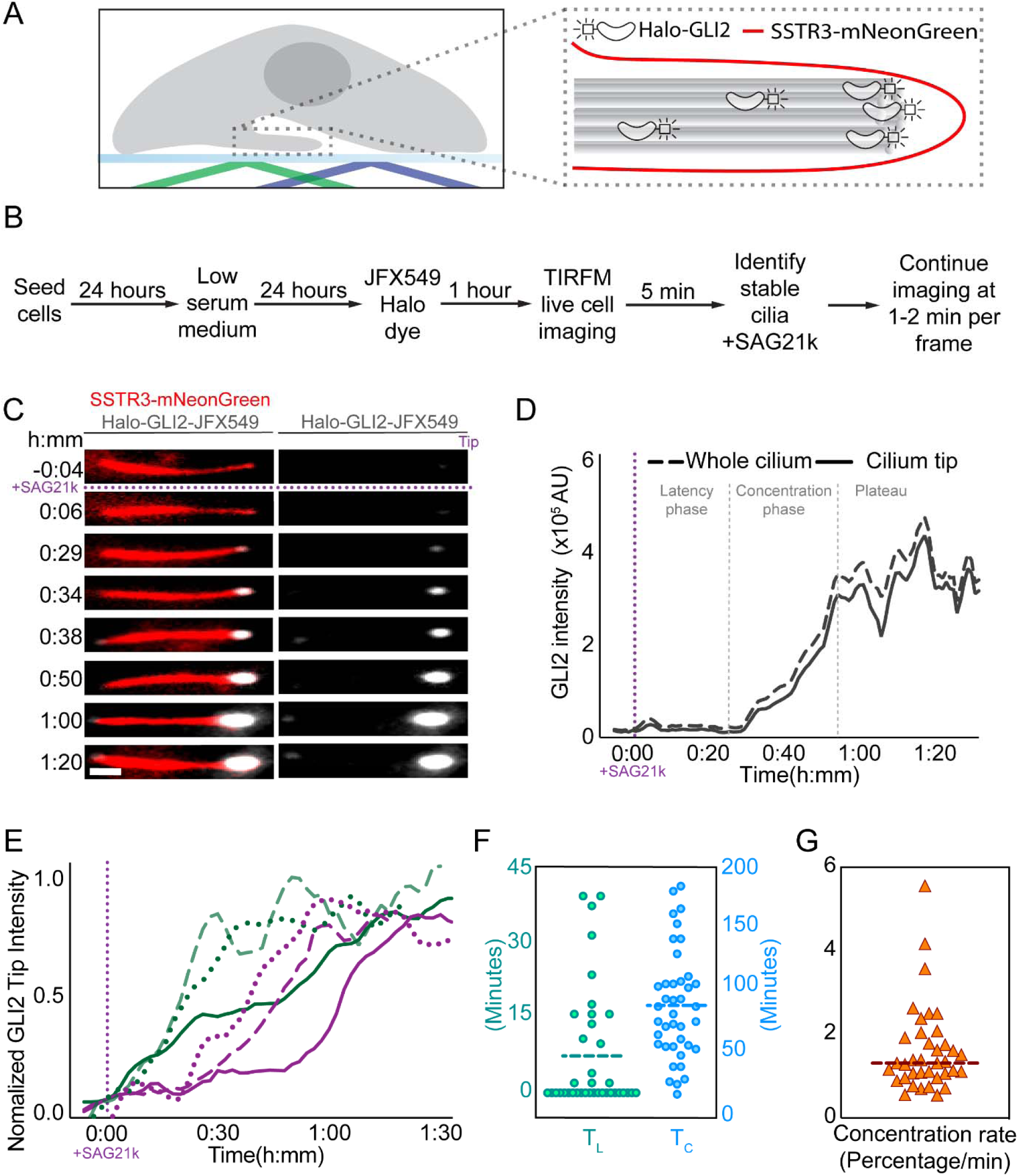
Real-time visualization of GLI2 accumulation at the cilium tip. (A) Schematic representation of a cell with the cilium parallel to the coverslip surface for TIRFM imaging (left). A close-up view of the cilium (right) shows the labeling of cilia by the membrane protein SSTR3-mNeonGreen (red) and Halo-GLI2 (gray). (B) Flowchart outlining the imaging protocol. (C) Representative montages from live-cell imaging of a cilium. Merged images display cilia in red and Halo-GLI2-JFX549 in gray (left). The individual Halo-GLI2-JFX549 channel is also displayed (right). Hh pathway activation occurred at t = 0:00 (h:mm) by the addition of SAG21k. The montages illustrate a relatively low Halo-GLI2-JFX549 signal at the cilium tip at the start of the experiment. After SAG21k addition, the Halo-GLI2-JFX549 signal at the tip gradually increased over time (Scale bar = 1 μm). (D) Intensity profiles of Halo-GLI2-JFX549 before and after SAG21k activation for the cilium in (C) The solid black line represents Halo-GLI2-JFX549 intensity at the cilium tip, demonstrating a consistent increase after SAG21k activation until reaching a plateau (defined as concentration time, T_C_). The dashed line indicates Halo-GLI2-JFX549 intensity throughout the whole cilium, including the tip and the shaft. (E) Normalized intensity profiles of Halo-GLI2-JFX549 from six additional examples, showing cilia with latency phase (T_L_) of varying durations. Green lines (dotted, dashed, and solid) are examples of three cilia with very small or no latency phase. Magenta lines (dotted, dashed, and solid) are examples of three cilia with significant latency phase. (F) The scatterplot displays GLI2 latency time (T_L_) and concentration time (T_C_) with the average indicated by the dashed line. (G) The scatterplot shows the GLI2 concentration rate, indicating the rate of normalized Halo-GLI2-JFX549 intensity increase during the concentration phase, with the average of 1.62% per minute depicted by the dashed line.

**Supplementary Figure 1.**
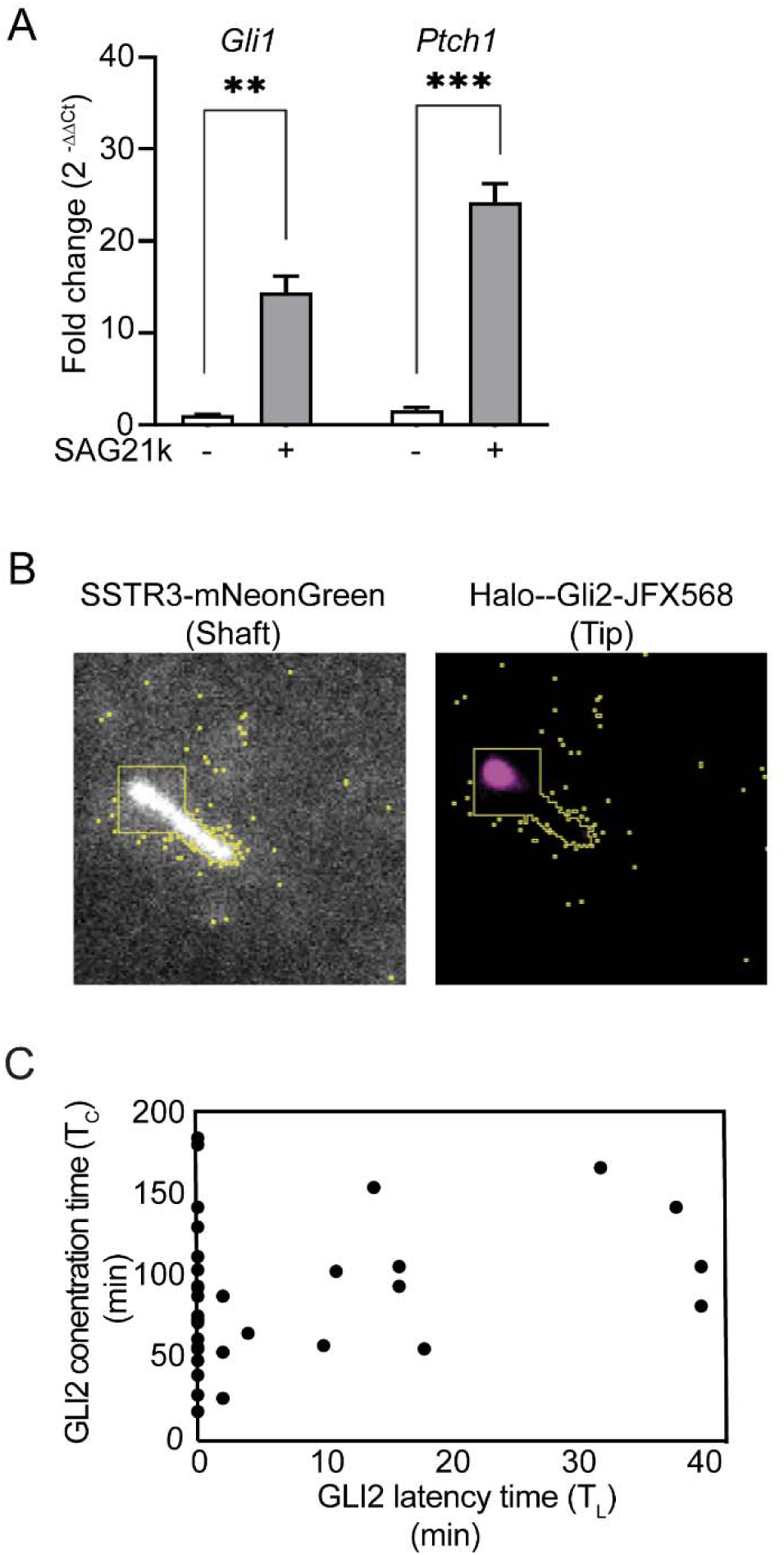
Real-time visualization of GLI2 accumulation at the cilium tip. (A) Bar graph showing the fold change in mRNA expression of *Gli1* and *Ptch1* by real-time PCR, with and without 12 hours of SAG21k treatment in NIH3T3 cells expressing Halo-GLI2. Each bar represents the mean of three independent experiments and the error bars represent standard error of the mean (SEM). Comparisons were carried out using a two-tailed t-test (p-value < 0.05). (B) A 25 x 25-pixel square mask created in the SSTR3-mNeonGreen channel was used to calculate the integrated GLI2 intensity in the Halo-GLI2-JFX549 channel. (C) Scatte r plot of TC versus TL of GLI2 in minutes.

Time-lapse images (Fig. 1C) showed an increase in Halo-GLI2-JFX549 fluorescence intensity over time in the cilium after SAG21k addition (Supplementary video 1). We quantitatively analyzed the integrated fluorescence intensity of GLI2 at the tip and the whole cilium (Fig. 1D). The representative intensity versus time plots for multiple cilia in Fig. 1E demonstrate that the fluorescence intensity profiles at the tip closely match those of the whole cilium (Supplementary Fig. 1B; note that fluorescence intensity at the base is not reliably captured in TIRFM imaging, since the base is positioned at variable height away from the evanescent field created by TIRF illumination). This indicates that the localization of GLI2 at the cilium tip primarily results from the trafficking of new molecules into the cilium rather than the redistribution of existing molecules from the cilium shaft to the tip after pathway activation. The plot of GLI2 cilium tip intensity versus time revealed a consistent pattern characterized by three distinct phases (n=44; Fig. 1D). We define these as (1) Latency phase (T_L_), where GLI2 tip intensity shows minimal change compared to the tip intensity before SAG21k addition; (2) Concentration phase (T_C_), during which GLI2 tip intensity increases over time; and (3) Plateau phase, when GLI2 tip intensity reaches a maximum and remains at a relatively constant level. Quantitative analysis revealed that T_L_ varied from 0-40 minutes, with 59% of the cilia exhibiting no detectable latency time (green curves in Fig. 1E), and the average T_C_ was 88 ± 6.9 minutes (Fig. 1F-G). Additionally, the lack of correlation between T_L_ and T_C_ suggests that the mechanisms determining GLI2 entry into the cilium and its transport and tip enrichment are distinct (Supplementary Fig. 1C).

In summary, we developed a TIRFM assay to image the dynamics of the primary Hh transducing transcription factor, GLI2. Our findings indicate that GLI2 molecules enter the cilium within 40 minutes of SAG21k addition, and accumulation at the cilium tip compartment peaks within two hours of pathway activation. Thus, the enrichment of GLI2 at the cilium tip compartment is an early event in Hh signaling.

### GLI2 is an IFT cargo

To further investigate the mechanism by which GLI2 concentrates at the cilium tip, we focused on the initial 2-hour time window following Hh pathway activation. We combined fast-imaging acquisition rate of 200 ms/frame with high laser power, which permitted approximately 3 minutes of imaging per cilium before photobleaching occurred. Due to the variability in GLI2 accumulation kinetics between cells (Fig. 1E-1G), we scanned for cilia with discernable signals in the shaft and selected them for imaging, thereby excluding those in the latency or plateau phase. This protocol (see Methods) enabled the visualization of Halo-GLI2-JFX549 at single particle resolution in the cilium shaft.

Visual examination of the movies from these experiments clearly showed GLI2 particles moving both towards and away from the cilium tip (Fig. 2A and Supplementary video 2). Kymographs were generated from line region of interest (ROI) drawn along the ciliary shaft in time-lapse movies (highlighted by dotted white boxes in Fig. 2B) to exclude the saturated signal at the cilium tip. Kymograph analysis revealed extensive bidirectional trafficking of GLI2, which was highly reminiscent of the IFT trains within the cilium (Fig. 2B). To determine whether GLI2 is associated with the IFT system, we created an NIH3T3 cell line that stably co-expresses both Halo-GLI2 and mNeonGreen-IFT88. We co-imaged the trafficking of Halo-GLI2-JFX549 and mNeonGreen-IFT88 nearly simultaneously at a frame rate of 700 ms/frame. Kymographs clearly showed overlapping co-trafficking tracks of GLI2 and IFT88 particles in both the anterograde and retrograde directions (marked in brown in Fig. 2C, Supplementary video 3).

**Figure 2.**
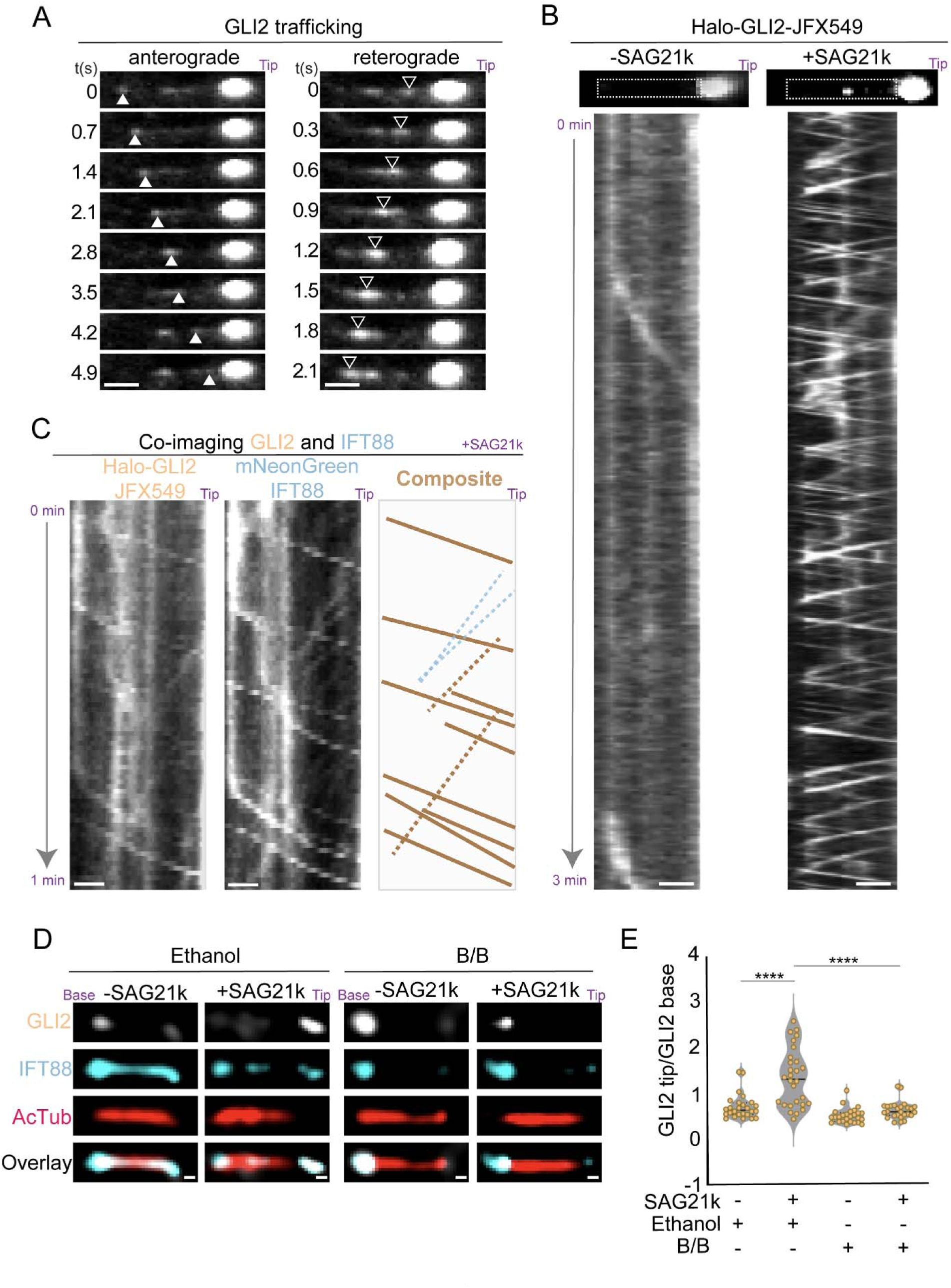
GLI2 is an IFT cargo. (A) Montages of Halo-GLI2-JFX549 trafficking in the cilium 17 minutes after the addition of SAG21k highlighting anterograde (solid arrowheads; left) and retrograde (open arrowheads; right) movement. (B) Kymographs of Halo-GLI2-JFX549 live imaging under two conditions: before SAG21k addition (-SAG21k, left) and 17 minutes after SAG21k addition (right). The image above each kymograph shows the corresponding cilium. The white dashed box on each image represents the area used to generate kymographs, excluding the saturated tip signal. Scale bars = 1μm. Each kymograph represents a time duration of 3 minutes. (C) Kymographs from the co-imaging of Halo-GLI2-JFX549 and mNeonGreen-IFT88 in the same cilium 18 minutes after SAG21k addition. The composite shows an overlay of individual channels. Co-transport of GLI2 and IFT is indicated by brown tracks in the anterograde direction and brown dashed tracks in the retrograde direction. Blue dashed lines show retrograde IFT tracks with no corresponding GLI2 track. (D) Representative images from the immunofluorescence assay of the i3A3B cells treated with 1% ethanol or 50 nM B/B homodimerizer in the absence or presence of SAG21k. Scale bars = 1 μm. (E) Violin plot showing the ratio of GLI2 intensity at the cilium tip to the cilia base. The value of n represents >26 cilia in each condition over three independent experiments. Statistical significance was determined using a two-tailed t-test (p-value < 0.0001).

We examined the impact of anterograde IFT inhibition on GLI2 trafficking and tip localization using a previously described system for acute kinesin-2 (KIF3A/KIF3B) inhibition^62^. For this, we utilized a Kif3a-/-Kif3b-/- NIH3T3 Flp-In cell line that stably expresses N-terminally DmrB-tagged KIF3A and KIF3B motors that are inhibited by the addition of B/B homodimerizer^62^. To verify the cell line expressing inhibitable KIF3A/KIF3B (referred to as i3A3B) motors, we incubated the cells with B/B homodimerizer for 8 hours and found that this treatment resulted in the complete loss of primary cilia as expected^62^ (Supplementary Fig. 2B). Next, we performed a shorter 30-minute B/B homodimerizer treatment to inhibit Kif3a/Kif3b while preserving the ciliary architecture and assessed GLI2 localization at the cilium tip via immunofluorescence. We found that under this condition, nearly 70% of cilia maintained standard lengths. Cilia exhibiting aberrant morphology were excluded from further analysis (see Methods). In control experiments, 62% of cells treated with SAG21k + DMSO for 30 mins showed GLI2 fluorescence signal above the background at the cilium tip. In contrast, in cells treated with SAG21k + B/B homodimerizer for 30 minutes, only 7% of cells had discernable GLI2 signal at the cilium tip (Fig. 2D and Supplementary Fig. 2A). Interestingly, in the B/B homodimerizer-treated cells, GLI2 localized at the ciliary base with IFT (Fig. 2D). Analysis of GLI2 intensity ratio between the cilium tip and base showed that the tip-to-base ratio is less than 1 in the B/B homodimer condition, and greater than 1 in control cells treated with SAG21k (Fig. 2E).

**Supplementary Figure 2.**
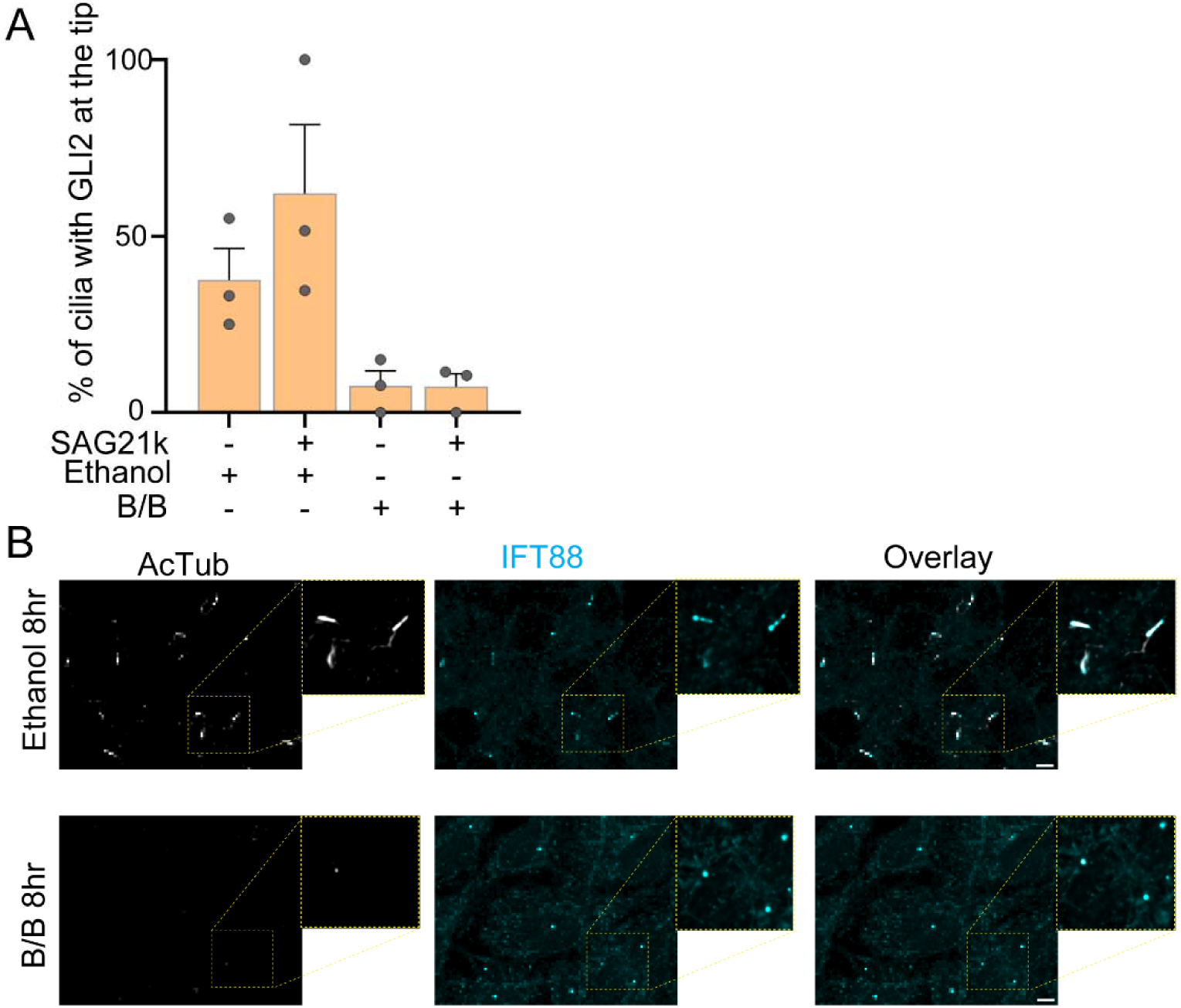
GLI2 is IFT-associated. (A) Bar graph showing the percentage of primary cilia showing GLI2 located at the tip under each condition. The gray points indicate mean values from three separate experiments. Error bars represent the standard error of mean (SEM). (B) Representative images of NIH3T3 Kif3a-/- kif3b-/- cells expressing inhibitable Kif3A and Kif3B, showing the presence of cilia upon ethanol treatment for 8 hours and their absence upon treatment with the B/B homodimerizer for 8 hours. Samples were stained with IFT88 and acetylated-α-tubulin antibodies to mark the cilia. The scale bars represent 5 μm.

Collectively, these experiments reveal that GLI2 is a bona fide cargo of IFT trains, conclusively establishing a transport role for IFT in Hh signaling. Perturbing IFT transport affects GLI2 localization at the tip compartment during the early time window after pathway activation. Thus, IFT transport is likely the predominant mechanism of GLI2 transport for its accumulation at the cilium tip compartment.

### GLI2 loading on IFT trains is temporally regulated during Hedgehog signaling

We analyzed the timing of GLI2 loading onto IFT trains and whether they were preferentially loaded onto anterograde IFT trains during the concentration phase of GLI2 accumulation at the cilium tip. We used kymographs from fast-acquisition single-particle imaging (Fig. 2) to quantify the frequencies and directionality of GLI2 and IFT88 track in each cilium before and at various time points after SAG21k addition. Each cilium was imaged for 3-5 minutes at a specific time point after pathway activation, and data were binned into 15-minute intervals to examine changes in GLI2 and IFT track frequencies over time.

In the absence of SAG21k, we detected very few moving GLI2 particles (Fig. 3A), even though significant IFT trafficking was evident under the same condition (Fig. 3B). Pathway activation triggered a burst of GLI2 movement (Fig. 3A). Binned time-interval analysis revealed an increased frequency of GLI2 movement during the first 45 minutes after SAG21k addition (Fig. 3A bins 0-15, 16-30, 31-45). The movement frequency reduces at later intervals, particularly noticeable after 60 minutes. The analysis revealed that the initial GLI2 movement was significantly biased in the anterograde direction (Fig. 3A). In contrast, a similar analysis of IFT88 particles showed no significant changes in response to SAG21k activation or time-dependence variation post-SAG21k addition. (Fig. 3B). While IFT88 also exhibited more anterograde than retrograde tracks, the directional bias was lower compared to GLI2 (Fig. 3B). These data suggested that anterograde IFT trains may preferentially load GLI2 as cargo following pathway activation. To examine this further, we calculated the percentage of IFT88 tracks that corresponded with GLI2 tracks across different time bins relative to SAG21k addition (Fig. 3C). In the absence of SAG21k, no GLI2 association with IFT trains was detected. This association increased to approximately 60% at 30-45 minutes post-SAG21k addition, then decreased to below 10% after 60 minutes. Given that the labeling efficiency of Halo tags with JFX549 ligand is expected to be less than 100%, the reported percentages likely represent the lower limit of the loading of GLI2 on IFT trains.

**Figure 3.**
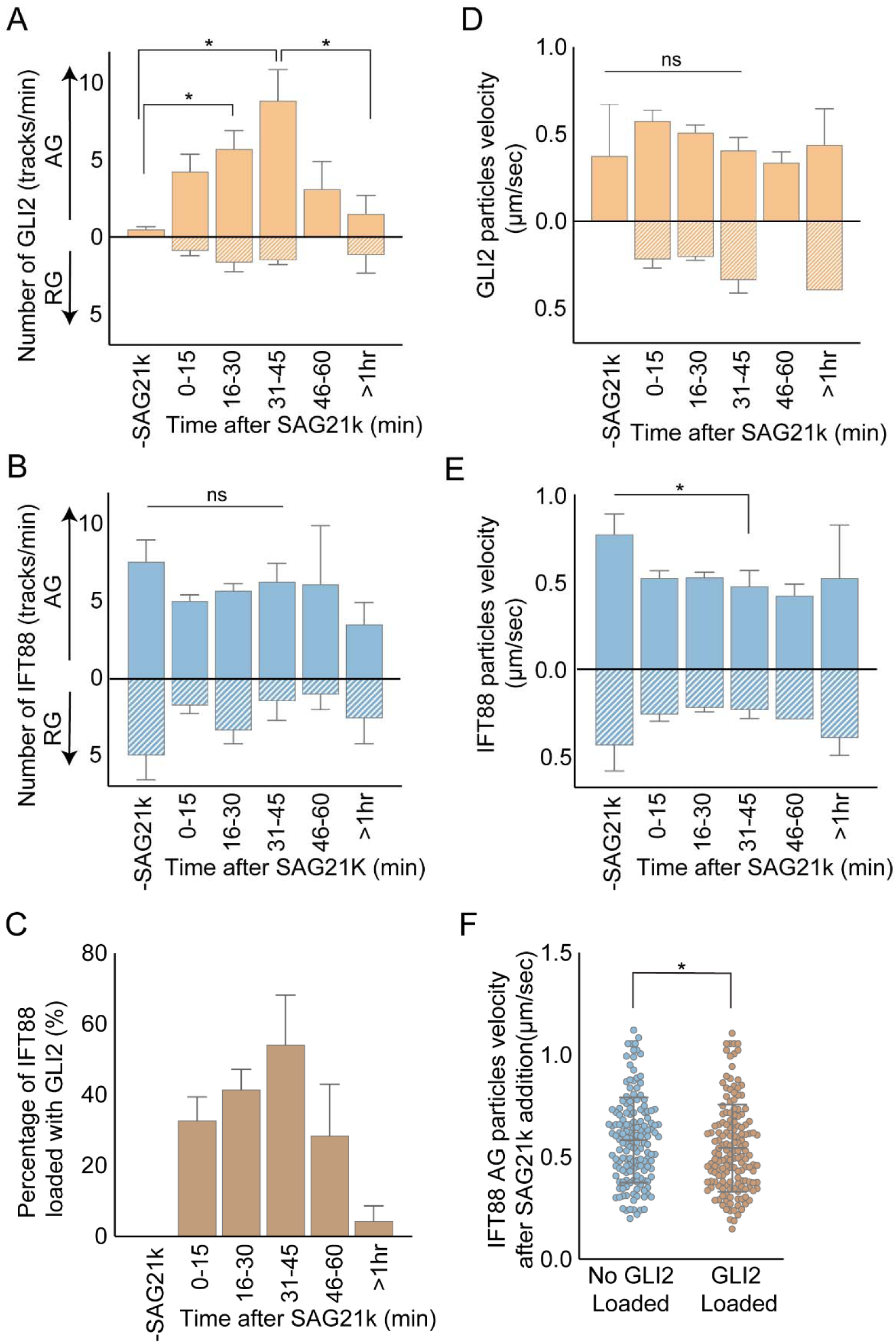
GLI2 association with IFT trains is temporally regulated during Hedgehog signaling. Bar graph showing (A) the number of GLI2 tracks per minute (n=37 cilia) and (B) the number of IFT88 tracks per minute (n=28 cilia) before and after the addition of SAG21k. Anterograde (AG) and retrograde (RG) datasets are shown on the same graphs, with AG above the 0 mark and RG below the 0 mark, respectively. (C) Bar graph showing the percentage of IFT88 tracks loaded with GLI2 (n=28 cilia). (D) Bar graph presenting the velocity in μm/sec of GLI2 particles and (E) IFT88 particles. AG and RG velocity values are above and below the 0 mark, respectively. The error bars represent SEM. (F) The scatter plot shows the velocities of IFT88 particles with and without observable GLI2 signal (n=28 cilia). Mean velocity with standard deviation is shown in each case. Statistical significance was determined using a one-tailed t-test (p-value < 0.05). *ns* indicates non-significant results.

We next examined the velocities of the moving GLI2 and IFT particles. The velocities of GLI2 particles were 0.5658 ± 0.01393 μm/sec in anterograde and 0.2695 ± 0.03087 μm/sec in the retrograde direction. IFT88 particles moved with a velocity of 0.5456 ± 0.01385 μm/sec in the anterograde direction and 0.2985 ± 0.01573 μm/sec in the retrograde direction (Supplementary Fig. 3A). This is consistent with reported IFT movement velocities^63^. No significant change in GLI2 particle velocity was observed before and after SAG21k addition (Fig. 3D). However, a ∼30% decrease in IFT velocity was noted following pathway activation (Fig. 3E). This reduction also corresponds to a slight decrease in anterograde velocity for GLI2-loaded IFT tracks compared to unloaded IFT tracks in the dual-imaging experiment (Fig. 3F).

**Supplementary Figure 3.**
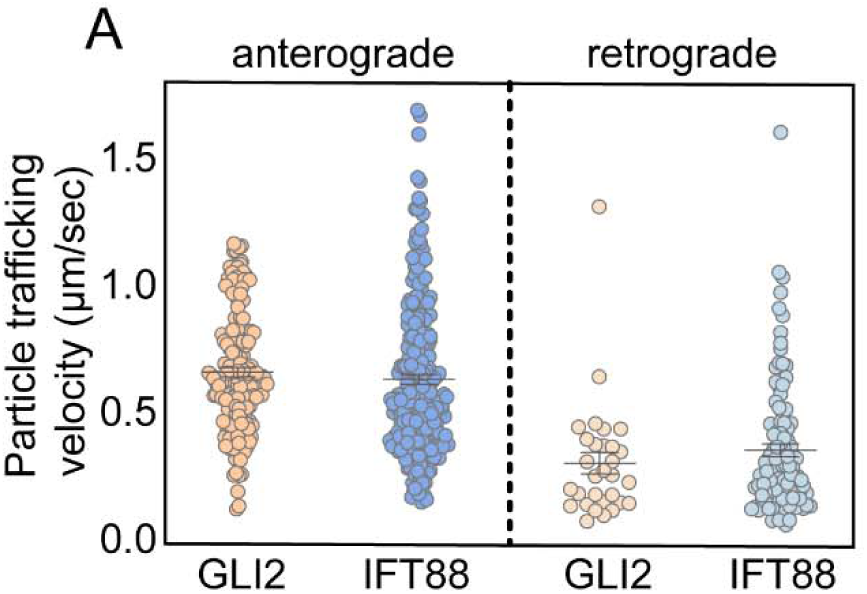
GLI2 association with IFT trains is temporally regulated during Hedgehog signaling. The scatter plot shows the anterograde and retrograde velocities of GLI2 and IFT88 particles.

Taken together, these data demonstrate that GLI2 loading on IFT is induced by Hh pathway activation, is temporally tightly regulated, and is initially biased towards the anterograde direction after pathway activation, leading to the formation of a GLI2-enriched tip compartment.

### Synchronized IFT-dependent cilium tip accumulation of KIF7 and GLI2 after pathway activation

The non-motile ciliary kinesin KIF7 co-localizes with GLI2 at the cilium tip^23,27,28^. To investigate the relative kinetics of KIF7 and GLI2 tip accumulation, we generated an NIH3T3 cell line co-expressing KIF7-Snap (using an inducible lentivirus system) and Halo-GLI2 (using a Flp-In^TM^ expression system) and simultaneously imaged the localization of both proteins at the distal cilium tip. KIF7 and GLI2 were labeled with Snap-Star505 and Halo-JFX549 dyes, respectively, while the cilium was marked with SiR-Tubulin. The co-imaging results revealed a simultaneous increase in intensity for both KIF7 and GLI2 at the cilium tip (Fig. 4A). In the representative image, the quantitative intensity profile at the cilium tip for both proteins entered the concentration and plateau phase in parallel, with KIF7 and GLI2 saturating at approximately T_C_=35 minutes (Fig. 4B, top panel). Correlation analysis of KIF7 and GLI2 intensity profiles over time revealed a high correlation during the concentration phase (Fig. 4B bottom panel, highlighted in pink). The mean correlation coefficient (CC) from all the concentrate phases of the cilia is 0.76 (n=9) (inset in Fig. 4B). This observation suggests a coordinated and temporally synchronized accumulation of KIF7 and GLI2 at the tips of the primary cilium.

**Figure 4.**
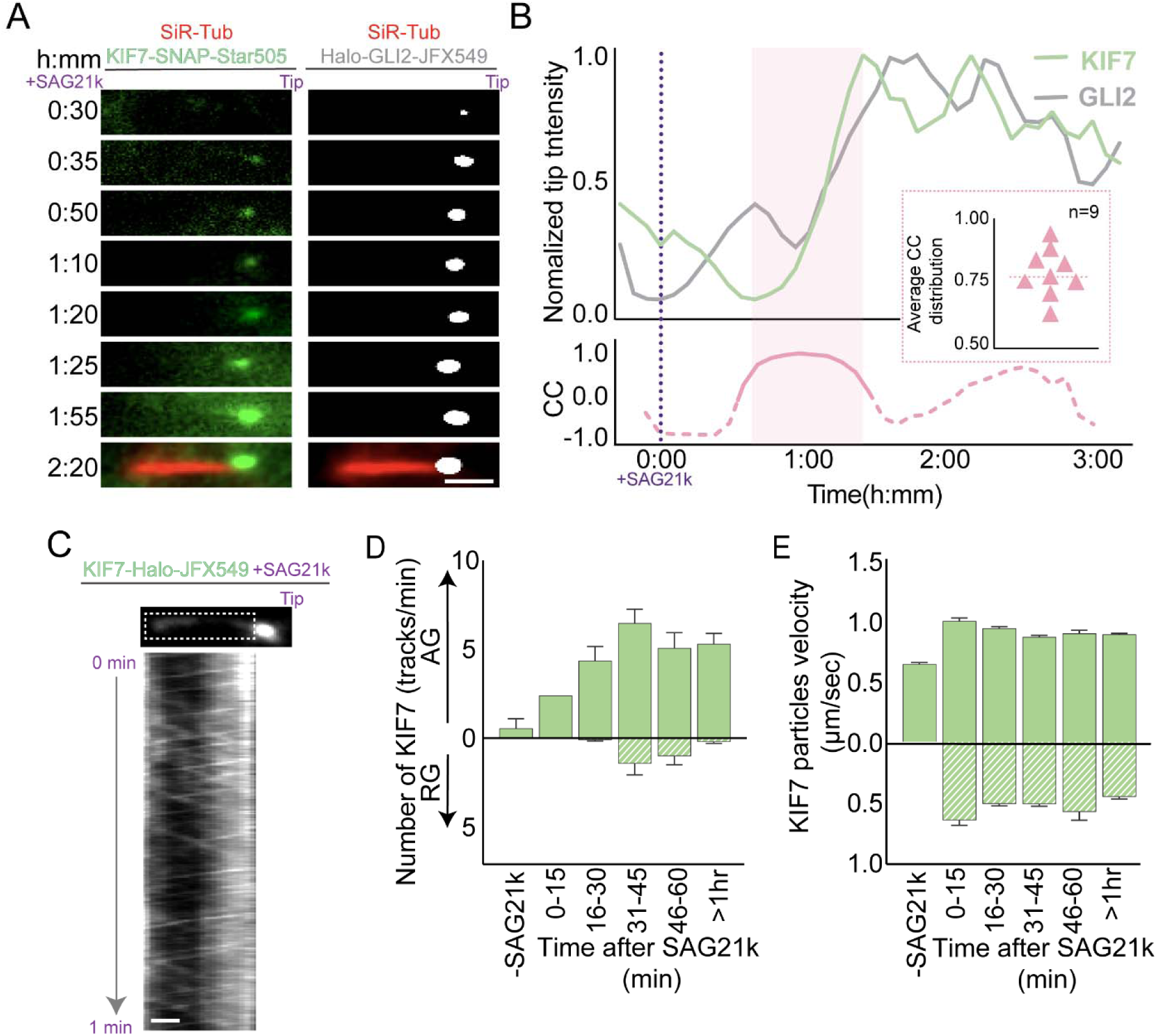
Synchronized cilium tip accumulation of KIF7 and GLI2 after pathway activation is IFT-dependent. (A) Montages from co-imaging of KIF7-Snap-Star505 (green) and Halo-GLI2-JFX549 (gray) in the same cilia (marked in red with SiR-Tubulin). 1 μM SAG21k was added at t = 0:00 (h:mm). Scale bar = 1 μm. (B) Normalized tip intensity of KIF7 and GLI2 corresponding to the cilia displayed in (A) plotted as a function of time. The bottom graph shows Pearson’s correlation coefficient change as a function of time after pathway activation for the same cilia. The inset shows the correlation coefficient calculated from the co-imaging dataset (n = 9 cilia). The mean correlation coefficient was 0.76, indicated by the dashed line. (C) Kymograph showing the bidirectional IFT-dependent trafficking of KIF7. Scale bar = 1 μm. (D) Bar graph showing the number of KIF7 tracks per minute (n = 62 cilia) and (E) The velocity of KIF7 particles in μm/sec (n = 62 cilia). AG and RG datasets are above and below the 0 mark, respectively. Error bar represents SEM values in each condition.

Based on the results from the KIF7 and GLI2 co-imaging assay and prior work in fixed cells^62,64^, we hypothesized that KIF7 is likely to be an IFT cargo. To test this, we generated an NIH3T3 cell line stably expressing KIF7-Halo (Flp-In^TM^ expression system). The Hh responsiveness of this cell line was tested by real-time PCR of the mRNA level expression of *Gli1* and *Ptch1* (Supplementary Fig. 4). Fast acquisition imaging of KIF7-Halo (200 ms/frame) showed IFT-like movement in the cilium (Supplementary video 4), with kymographs clearly showing bidirectional trafficking (Fig. 4C). Quantitative analysis of kymographs showed that KIF7, like GLI2, is an IFT cargo. Upon Hh pathway activation with SAG21k, we observed a burst of KIF7-loaded anterograde IFT movement, resulting in KIF7 tip accumulation (Fig. 4D-E).

**Supplementary Figure 4.**
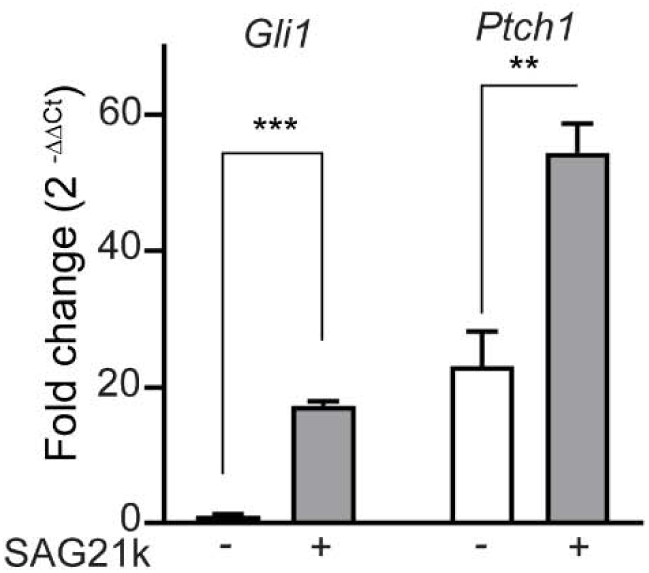
Synchronized IFT-dependent cilium tip accumulation of KIF7 and GLI2 after pathway activation. The bar graph shows the fold change in mRNA expression of *Gli1* and *Ptch1* by real-time PCR, with and without 12 hours of SAG21k treatment in NIH3T3 cells expressing KIF7-Halo.

These data support a mechanism by which KIF7 and GLI2, core components of the Hh signaling complex, utilize IFT trains for localization to the cilium tip compartment.

### KIF7 is a gatekeeper for GLI2 entry into the primary cilium

Previous immunofluorescence studies have shown that mutations or deletion of KIF7 lead to the mislocalization of GLI2, causing it to spread along the ciliary shaft instead of concentrating at the cilium tip^55^. To investigate this further, we developed a CRISPR-Cas9 knock-out of KIF7 in NIH3T3 cells (Supplementary Fig. 5A and 5B). We quantified the levels of endogenous GLI2 in the cilium of Kif7-/- cells and compared them to WT cells both before and after SAG21k treatment for 1 hour in immunofluorescence assays. We found that (1) before SAG21k addition, there are significantly higher levels of endogenous GLI2 in KIF7-/- cilia compared to WT cilia. (2) After SAG21k addition, the increase in ciliary GLI2 levels is lower in KIF7-/- cilia compared to WT cilia. (3) GLI2 is mislocalized along the ciliary shaft both before and after SAG21k treatment (see representative images in Fig. 5A and quantification in Fig. 5B), consistent with prior report^55^. Collectively, these data suggested that KIF7 regulates the levels and localization of GLI2 both before and after pathway activation.

**Figure 5.**
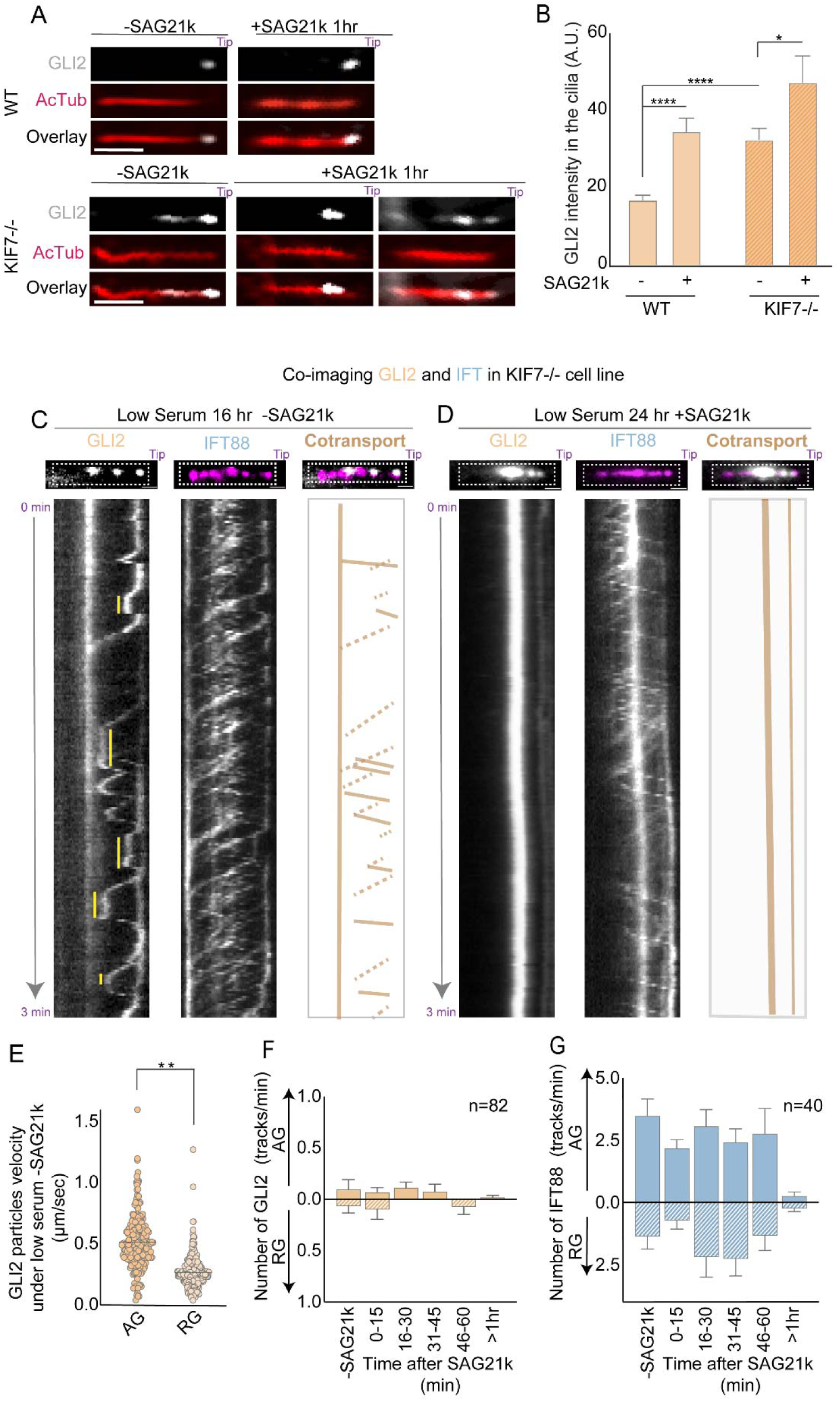
KIF7 regulates GLI2 entry into the primary cilium. (A) Representative images from an immunofluorescence assay in NIH3T3 WT cells and NIH3T3 KIF7 -/- cells stained for GLI2 and acetylated α-tubulin (cilia marker). Cells were treated with or without SAG21k for 1 hour. (B) Bar graph showing the intensity of GLI2 at the tip, quantified from the immunofluorescence datasets in (A) (n=84 cilia). Statistical significance was determined using a two-tailed t-test (p-value < 0.05). (C) Kymographs from co-imaging GLI2 and IFT88 in the NIH3T3 KIF7 -/- cilia under low serum for 16 hours without SAG21k and (D) under low serum for 24 hours after SAG21k activation. Yellow solid lines on the kymograph indicate pause events. Co-transport is shown by brown lines in the anterograde direction and brown dashed lines in the retrograde direction. Thick brown vertical lines show stationary particles. (E) Scatter plot showing AG and RT velocities of GLI2 particles under serum starvation of <16 hours without SAG21k (N =26 cilia). Comparisons were made using a two-tailed t-test with a p-value < 0.05. (F) Bar graph showing the number of tracks per minute for GLI2 and (G) IFT88 under low serum conditions for 24 hours followed by SAG21k treatment. All error bars represent SEM.

**Supplementary Figure 5.**
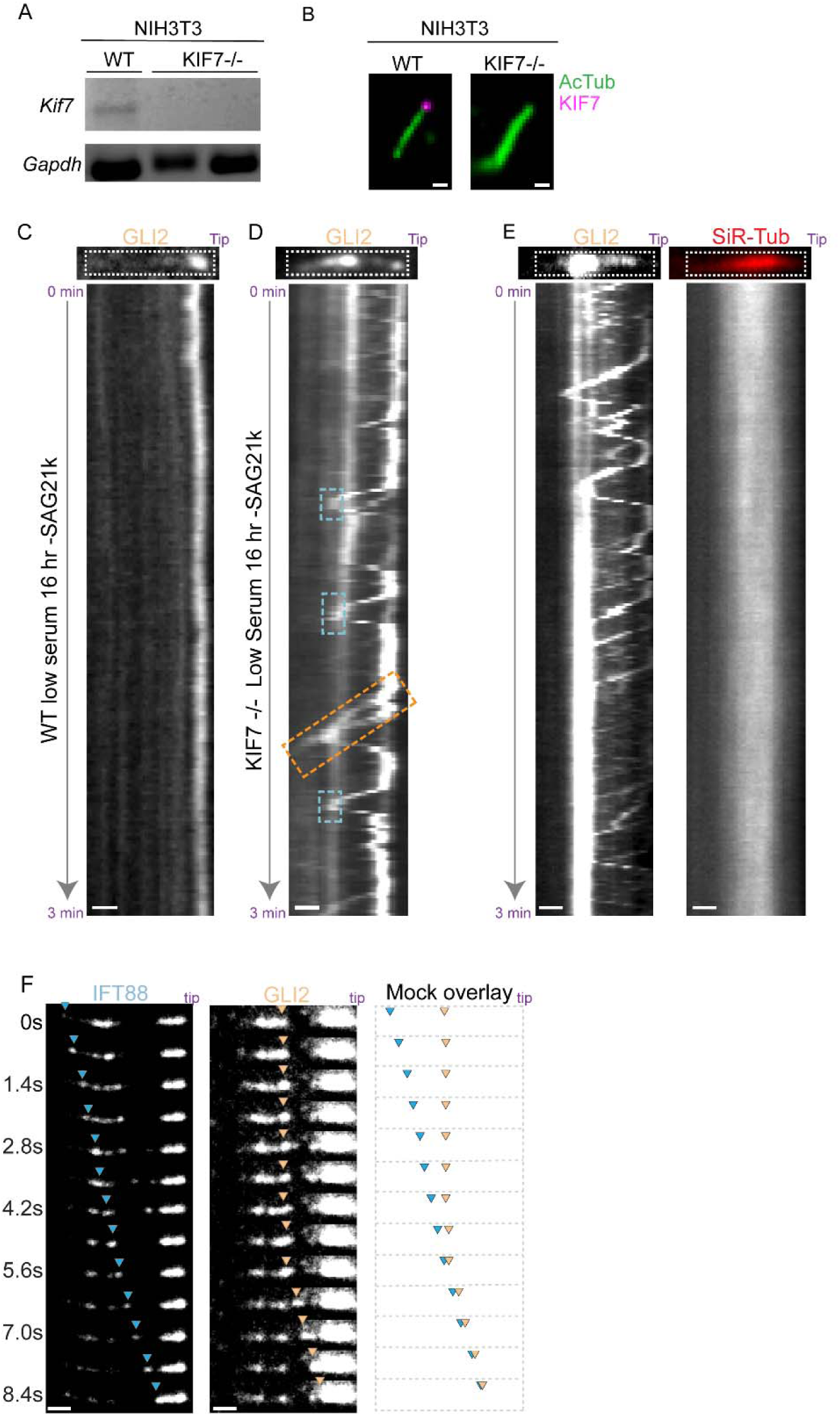
KIF7 is a gatekeeper regulating GLI2 entry into the primary cilium. (A) mRNA expression of *kif7* and *gapdh* in NIH3T3 WT and KIF7-/- cells. (B) Representative fixed images stained with anti-acetylated α-tubulin and KIF7 in WT and KIF7-/- cilia. (C) Kymograph of GLI2 imaging in WT cilia under low serum (without SAG21k). (D) Kymograph of GLI2 imaging in KIF7-/- cilia under low serum (without SAG21k), with the orange dashed box highlighting the complete removal of GLI2 from the cilium. The blue dashed boxes highlight multiple retrograde tracks that appear to pause or switch to an anterograde track. (E) Kymograph of GLI2 and SiR-Tubulin in KIF7-/- cilia serum-starved for 16 hours without SAG21k. (F) Montages of IFT88 and GLI2 in WT cells showing shaft clearance of GLI2 by IFT trains. The blue arrowhead follows the IFT particle, and the orange arrowhead follows the GLI2 particle. Scale bar = 1 μm.

We investigated KIF7’s role in regulating GLI2 localization in the cilium in the absence of pathway activation. We performed live imaging of Halo-GLI2-JFX549 and mNeonGreen-IFT88 in KIF7 -/- and WT cilia without SAG21k addition. For these experiments, we performed 16 hours of serum starvation to promote ciliogenesis. In WT cilia, no significant accumulation or movement of GLI2 was observed in the absence of SAG21k (Supplementary Fig. 5C). In contrast, KIF7-/- cilia displayed GLI2 puncta in the ciliary shaft, consistent with immunofluorescence results (Fig. 5A). Live cell imaging revealed movement of GLI2 puncta in the cilium shaft in both anterograde and retrograde directions with velocities consistent with IFT-based transport. (Fig. 5C, left panel, and Fig. 5E), Co-imaging with IFT showed that GLI2 movement coincided with IFT tracks (Fig. 5C, Supplementary video 5, Fig. 5D). These data show that GLI2 is loaded onto IFT in the absence of KIF7 even without pathway stimulation by SAG21k. The absence of KIF7 additionally causes GLI2 to aberrantly detach from IFT within the cilium (Fig. 5C), in contrast to unloading at the cilium tip or base as observed in WT cells.

Interestingly, GLI2 puncta that reach the tip in KIF7-/- cilia do not stably associate with the tip (Fig. 5C, GLI2 channel). These observations contrast with WT cells where GLI2 molecules are stably associated at the distal tip with low levels of anterograde transport (Supplementary Fig. 5C). Instead, in the KIF7-/- cells GLI2 molecules at the cilium tip associate with retrograde IFT and are transported towards the base of the cilium. Occasionally, these puncta are observed to exit the ciliary shaft (Supplementary Fig. 5D, orange dashed box shows an example of cilium exiting GLI2 puncta). In general, three patterns of GLI2 dynamics were observed: (1) retrograde IFT-bound GLI2 puncta switches to an anterograde train and is redirected toward the tip, (2) retrograde IFT-bound GLI2 puncta switches to a pause state without a corresponding pause of the IFT train (yellow solid lines on GLI2 kymograph in Fig. 5C indicate pause events), and (3) sometimes GLI puncta can load onto an anterograde train after a pause and be re-delivered to the tip (examples in the blue dashed boxes in Supplementary Fig. 5D). Over time (such as at 24 hours after serum starvation), the paused assemblies of GLI2 accumulate additional protein, and become less dynamic (GLI2 kymograph Supplementary Fig. 4E).

To rule out the possibility that the positional changes in GLI arise from changes in the length of the cilium rather than switching on IFT trains, we monitored SiR-Tubulin intensity along the cilium. We found that tubulin intensity profiles along the cilia remain relatively stable over the time course of imaging (SiR-Tubulin channel in Supplementary Fig. 5E). Together with the observation that GLI2 puncta are associated with moving IFT trains, the data suggests that GLI2 dynamics are due to bidirectional trafficking rather than axoneme length fluctuations.

We next examined how GLI2 dynamics in KIF7-/- cilia change after pathway activation. Under these conditions, we observed larger puncta or patches of GLI2 within the ciliary shaft (kymographs in Fig. 5D; right panel). These GLI2 assemblies were mostly sedentary, with minimal movement of GLI2 molecules up to 2 hours after SAG21k addition (Fig. 5F). In these cilia, moving IFT trains were initially observed but these movements were greatly reduced after 1 hour of pathway activation, potentially because they are trapped within or blocked by the large GLI2 assemblies (Fig. 5G). In contrast, in WT cilia, we occasionally captured events where paused GLI2 puncta in the ciliary shaft were transported to the tip by a moving anterograde IFT train, thus clearing the shaft (Supplementary Fig. 5F).

These data indicate that GLI2 can load on IFT in the absence of KIF7. However, without KIF7, both the temporal regulation of GLI2 transport and its spatial restriction to the cilium tip are severely compromised. Thus, this non-motile kinesin emerges as a major regulator of GLI2 transport by the motile IFT transport system in the cilium.

## DISCUSSION

GLI2 accumulation at the distal cilium tip is important for accurate Hh signaling, contributing to precise transcriptional output in response to ligand. In this study, we defined the dynamics and mechanism of GLI2 transport by developing an assay for real-time cilium imaging of the transcription factor GLI2, the conserved pathway regulator KIF7, and the core ciliary IFT machinery. Our findings reveal how two ciliary cytoskeletal systems, one motile and one non-motile, synergize to establish a GLI2-enriched cilium tip compartment for Hh signal transduction.

We found that GLI2 (Fig. 2) and KIF7 (Fig. 4C) are transported to the cilium tip as IFT cargoes. While the IFT trains’ movement appears largely Hh-independent, GLI2 is loaded onto these trains within a restricted time window after pathway activation, with a striking increase in movement during the concentration phase (Fig. 3A, B and, Fig. 6). The movement largely ceases below detection levels within 1-2 hours after SAG21k addition in our assay. The overall kinetics of GLI2 tip accumulation observed here are consistent with fixed cell immunofluorescence data^21^ and proteomic analyses of ciliary GLI2 accumulation^65^. Importantly, IFT-mediated GLI2 transport during the concentration phase is significantly anterograde-biased, leading to cilium tip accumulation (Fig. 6). Future studies will investigate whether the cessation of GLI transport is due to negative regulation of IFT loading upon reaching a tip occupancy threshold or to the depletion of these proteins in the cytoplasmic space outside the cilium. Interestingly, albeit at a much lower frequency, we also observe GLI2’s association with retrograde IFT trains. This observation underscores the notion that GLI2 traffics through the cilium tip compartment en route to its final destination in the nucleus. The temporally and directionally regulated loading of Hh pathway proteins is reminiscent of the increased tubulin loading on IFT during ciliogenesis in Chlamydomonas^29^. Thus, the role of IFT as an omnipresent cilium shuttle combined with tightly regulated cargo loading for specific cellular processes, may represent a common theme in protein trafficking within the cilium.

**Figure 6.**
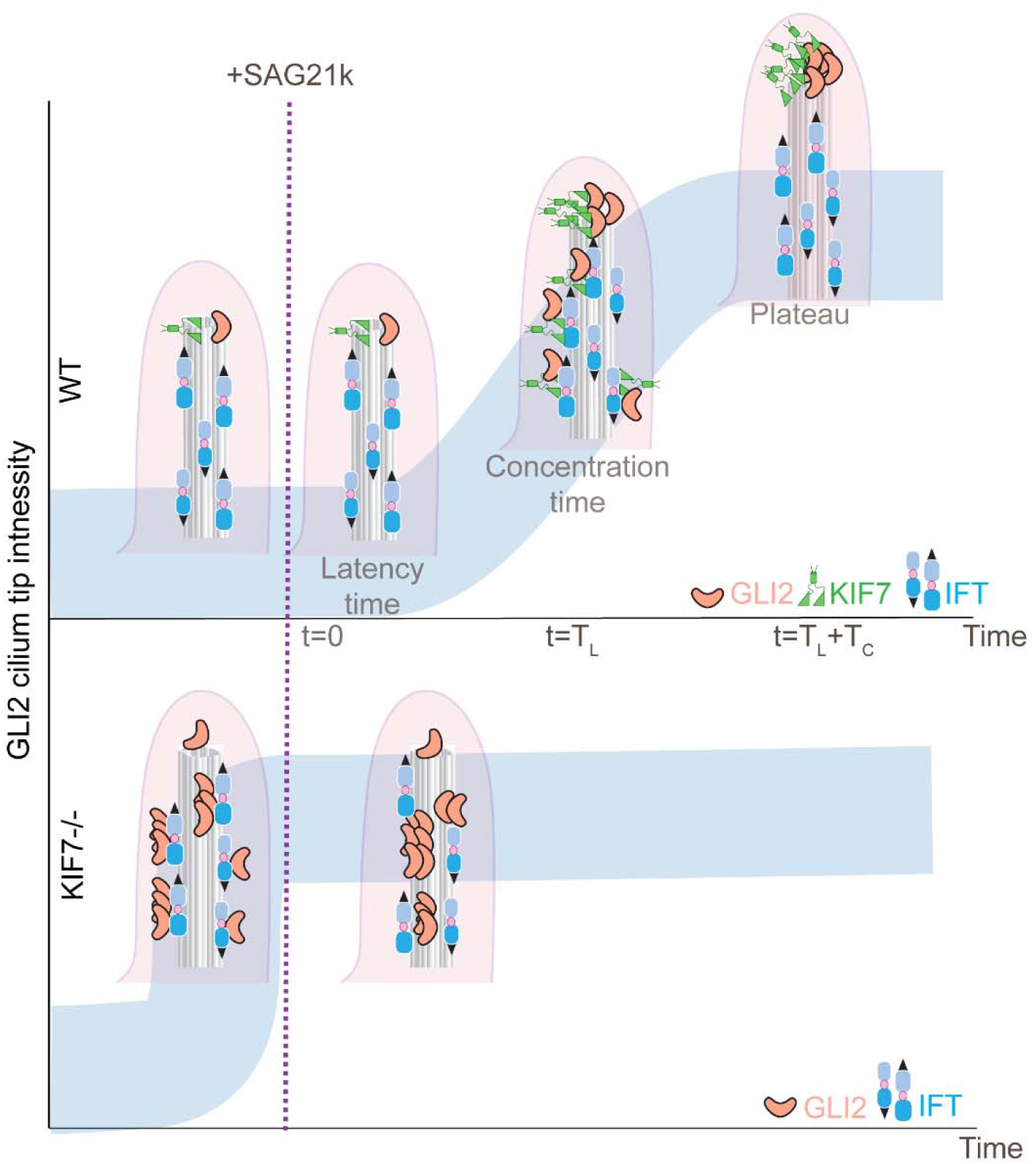
Schematic illustrating the ciliary dynamics of GLI2. The top panel depicts the WT condition, where the ciliary trafficking of GLI2 and KIF7 after pathway activation occurs in three phases: latency, concentration time, and plateau. The highest levels of trafficking of GLI2 and KIF7 to the cilium tip occur during the concentration phase. Both GLI2 and KIF7 are IFT cargoes transported along the cilium shaft. At the distal tip, the proteins are off-loaded from IFT, and KIF7 serves to tightly anchor GLI2, forming the cilium tip compartment. The bottom panel shows the KIF7-/- condition, where increased GLI2 entry is observed before pathway activation. GLI2 molecules arriving at the cilium tip are not retained and instead switch to the retrograde IFT trains. Moving GLI2 puncta are frequently deposited in the ciliary shaft, where they accumulate over time. The shaft accumulation persists even upon pathway activation, resulting in larger non-dynamic GLI2 assemblies, thus obstructing the formation of the concentrated cilium tip compartment of GLI2 seen in the presence of KIF7.

While KIF7 is well established as a conserved Hh pathway regulator, precisely defining its mechanistic role has been challenging due to its dual function as a modulator of cilium architecture^66^, as well as a regulator of both Hh signaling activation and repression^23,27,28,67^. We observe that the characteristics of KIF7 transport by IFT closely mirror those of GLI2 trafficking (Fig. 4A). Our findings illuminate three critical roles for KIF7 in Hh signaling (Fig. 6). First, quantitative analysis in KIF7-/- cells shows that high levels of GLI2 are present in the cilium shaft even before pathway activation^55^, suggesting that KIF7 potentially restricts Gli2 loading onto IFT trains before Hh activation (Fig. 5). These observations correlate with increased levels of GLI2-A in KIF7-/- cells when the pathway is repressed^23,55^. Second, in the absence of KIF7, GLI2 molecules are not tightly confined to the cilium tip. Instead, we observe a striking removal of tip-bound Gli2 by retrograde transport (Fig. 5C and Supplementary Fig. 5D and E). Thus, KIF7 is critical for stably tethering GLI2 at the cilium tip (Fig. 5). Third, in WT cilia, GLI2 molecules noticeably “stuck” in the ciliary shaft are cleared by anterograde IFT trains and deposited at the cilium tip (Supplementary Fig. 5F). However, in KIF7-depleted cells, GLI2 puncta trapped along the shaft are not cleared (Fig. 5D) The lack of clearance may be attributed to two reasons. As previously hypothesized^50,55^, the absence of KIF7 might result in structural defects in the axoneme, creating sites for GLI2 accumulation. Alternatively, KIF7 might promote GLI2 transport by increasing its association with anterograde IFT. Regardless, the absence of KIF7 results in the trapping of GLI2 molecules along the axoneme which might be the reason for the reduction in GLI2-A observed after pathway activation in KIF7-/- cells (Fig. 6)^23,27,28,56^. Our findings suggest that KIF7 acts as a gatekeeper for GLI2, regulating its entry into the cilium before Hh activation and its exit from the cilium tip after Hh activation. Consequently, this non-motile kinesin assumes an unconventional role as a cargo regulator in cilium-dependent Hh signaling.

How are GLI2 and KIF7 linked to IFT trains? While defining the precise molecular connections within the KIF7-GLI2-IFT network requires additional studies, our findings, along with other reports, provide valuable insights. Our results suggest that KIF7 is not necessary for GLI2’s association with IFT trains, implying a direct interaction of GLI2 with one or more IFT components. Co-imaging experiments using labeled IFT88 and GLI2 rule out the possible mechanisms where GLI2 is transported by an alternative non-IFT motor or through an IFT-B independent KIF3A-KIF3B-KAP3 system^49^. Prior work suggests that IFT27 has a role in GLI2 and KIF7 tip localization^26,68^. However, in the absence of IFT27, GLI2 is present along the axoneme shaft, suggesting that IFT27 is not strictly required for GLI2 transport into the cilium^68^. We have previously shown that KIF7 directly binds GLI, and the localization of KIF7 at the cilium tip is dependent on GLI2 and GLI3^54^. Given that KIF7 is also an IFT cargo and that their cilium-tip accumulation kinetics are similar, it is likely that KIF7 and GLI remain in a complex during transport to the cilium tip. Whether KIF7 uses GLI as an adapter or has an independent binding site on IFT still remains unknown. In either case, GLI switches from using IFT for transport along axonemes to using KIF7 as a cilium tip localization scaffold. The binding of GLI2 to KIF7 increases the the anchoring of the Hedgehog Signaling Complex proteins at the cilium tip compartment. Future biochemical analyses will reveal how these Hh pathway proteins are linked to IFT trains and offloaded at the cilium tip compartment.

A unique feature of Hh signal transduction is the utilization of microtubule cytoskeleton as a scaffold for transcription factor regulation. In *Drosophila melanogaster*, the KIF7-homolog Costal-2, a non-motile kinesin-like protein, tethers the GLI-homolog Cubitus interruptus (Ci) to cytoplasmic microtubules^69^. Activation of the Hh pathway leads to Costal-2’s dissociation from microtubules and Ci’s entry into the nucleus. In cilium-dependent Hh signaling in vertebrates, the axoneme, rather than cytoplasmic microtubules, acts as the GLI-binding scaffold. The requirement for a distinct organelle with a highly specialized and polarized microtubule array necessitates additional regulatory steps for spatially and temporally restricted GLI localization to the cilium tip. Our findings suggest that this is achieved synergistically through an organelle-specific cytoskeletal system, IFT, and a pathway-specific cytoskeletal protein, KIF7, that collectively act to precisely position GLI2 at the cilium tip during vertebrate Hh signaling.

## Supporting information

Video corresponding to Fig 1C

Video corresponding to Fig 2A

Video corresponding to Fig 2C

Video corresponding to Fig 4C

Video corresponding to Fig 5C

## ACKNOWLEDGEMENTS

We thank Drs. Luke D. Lavis and Jonathan B. Grimm (Janelia) for a gift of JFX549-HaloTag ligand. We thank Dr. Robert E. Kingston for the EasyFusion Halo-mAID and the LentiCRISPR V2 plasmid, Dr. Peter Czarnecki for the pCDH-mNeonGreen-IFT88 plasmid, and Dr. Michael Blower for providing the doxycycline-inducible pLVX-TetOne-Puro vector and HEK239T cell line. We thank Dr. Mu He was feedback on the manuscript. R.S. was supported through the Pew Biomedical Scholars program, the American Cancer Society (Ellison Foundation Research Scholar) and NIH (NIGMS) 1R01GM145651; M.E. was funded by NIH (NIGMS) R35GM147641; D.M. was funded by NIH R01EY033035 and P30EY000331.

## Supplementary Videos

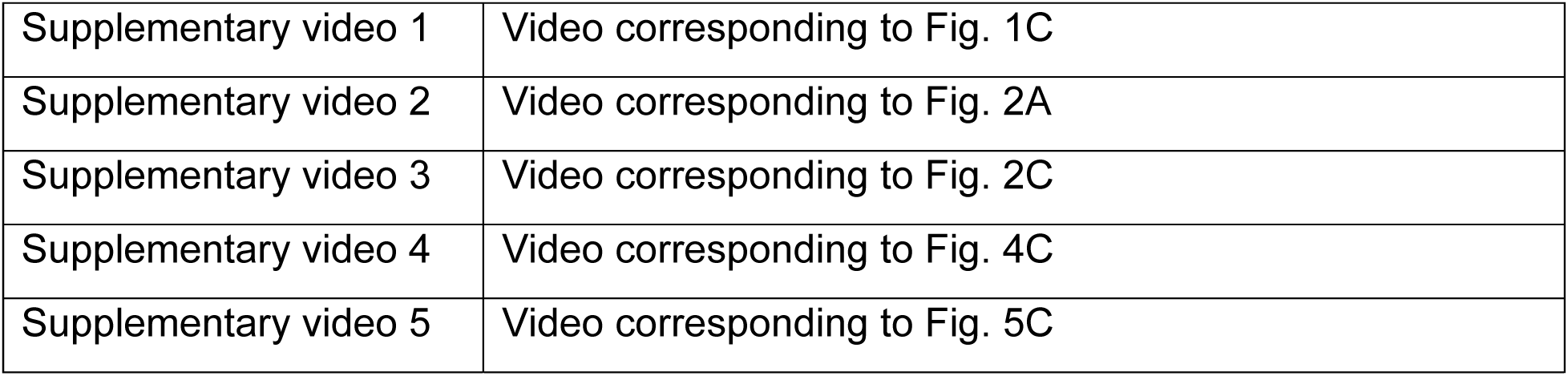

## MATERIALS and METHODS

### DNA constructs

The pEF5/FRT/DEST-3xHA/mGLI2 plasmid (Addgene: Plasmid #51250) was obtained from Addgene. The Halo-tag was amplified from EasyFusion Halo-Maid (a gift from Dr. Robert E. Kingston) and inserted at the N-terminus of mGLI2 within the pEF5/FRT/DEST-3xHA/mGLI2 plasmid using Gibson assembly. For long-duration imaging of KIF7, C-terminally tagged KIF7-mRuby3 and KIF7-Snap encoding cDNA were cloned into a doxycycline-inducible pLVX-TetOne-Puro vector (a gift from Dr. Michael Blower). For fast acquisition imaging of KIF7 movement in the cilium, KIF7-Halo cDNA was cloned into the pEF5/FRT/DEST vector. The pCDH-mNeonGreen-IFT88 plasmid was kindly provided by Dr. Peter Czarnecki. All DNA constructs were verified by full plasmid sequencing.

### Stable cell lines

#### NIH3T3 Halo-mGLI2

To develop a stably expressing Halo-mGLI2 cell line, an NIH3T3 host cell line containing a single copy of the Flippase Recombination Target (FRT) site was transfected with the pEF5-FRT-DEST-3XHA-Halo-mGLI2 plasmid and the Flp-ecombinase expression vector pOG44 (Life Technologies, Grand Island, NY). Transfection was performed using the JetPrime Polyplus transfection reagent (Polyplus, France). Following a 48-hour incubation, cells underwent selection using complete DMEM with 100 ug/ml of hygromycin. Single-cell isolation and expansion were conducted. The expression of Halo-GLI2 was verified via fluorescence microscopy.

#### NIH3T3 Halo-mGLI2 and KIF7-Snap

NIH3T3 Halo-GLI2 cells were infected with inducible lentiviral particles encapsulating KIF7-Snap-C-terminal. After 48 hours of infection, cells were selected using 2.5 ug/ml of puromycin combined with 100 ug/ml of hygromycin. A stable cell line was treated with 500 ng/ml doxycycline to induce KIF7-Snap expressing for 24 hours after seeding and prior to serum starvation.

#### NIH3T3 Halo-mGLI2 and mNeonGreen-IFT88

NIH3T3 Halo-GLI2 cells were infected with lentiviral particles encapsulating the pCDH-mNeonGreen-IFT88 plasmid. After 48 hours of infection, cells were selected using 500 ug/ml of geneticin combined with 100 ug/ml of hygromycin.

#### NIH3T3 mKIF7-Halo

NIH3T3 Flp-In cells were transfected with pEF5/FRT/DEST/mKIF7-Halo using the Flp-In^TM^ transfection protocol stated earlier. Selection started 48 hourspost-transfection using 100 ug/ml of hygromycin.

#### NIH3T3 KIF7-Halo and mNeonGreen-IFT88

The pCDH-mNeonGreen-IFT88 plasmid was used to infect KIF7-Halo-expressing cells lentivirus system. Selection was performed using 500 ug/ml of geneticin combined with 100 ug/ml of hygromycin.

#### NIH3T3 KIF7 -/-

The guide RNA sequences from Liu et al (Table 1) ^56^ wereused to generate the KIF7 knockout cell line by CRSPR-editing. Briefly, the cDNA corresponding to the guide RNA sequence was cloned into LentiCRISPR v2 plasmid (a kind gift from Dr. Robert E Kingston). Lentiviral soup containing LentiCRISPR v2 plasmid with KIF7 guide RNA sequence was used to infect NIH3T3 cells. 48 hours post-infection, the cells were selected using 10 ug/ml of blasticidin. After selection, the knockout was confirmed using immunofluorescence, mRNA amplification, and genomic DNA PCR

#### NIH3T3 Kif3a-/- kif3b-/- expressing i3A3B

The NIH3T3 Flp-In Kif3a-/- Kif3b-/- cell line was generated as described in Engelke et al. 2019 ^62^. The *iKif3a* and *iKif3b* sequences were cloned into the pUltra-hot plasmid (Addgene’s plasmid #24130), and together with pMD2.G (Addgene’s plasmid #12259) and psPAX2 (Addgene’s plasmid #12260), were transfected into HEK293T cells with jetPRIME to produce lentiviral particles. Three days after transfection, the supernatant containing the viral particles was collected and used to transduce the NIH3T3 Flp-In Kif3a-/- Kif3b-/- cells.

### Cell culture

HEK 293T cells were generously provided by Dr. Michael Blower’s Lab. The NIH3T3 cell line was purchased from BPS Bioscience (60409). Both cell lines were cultured in high-glucose Dulbecco’s Modified Eagle’s Medium (DMEM; ThermoFisher 11995081) supplemented with 10% HyClone Characterized Fetal Bovine Serum (FBS; Cytiva SH30071.03HI). Cultures were incubated at 37°C in a 5% CO2 and 95% relative humidity.

### Immunofluorescence

Cells were seeded on 35 mm plates with 18×18 mm coverslip. After an additional 24 hours of serum starvation in low-serum DMEM (0.2% FBS) containing 1 μM SAG21k (Fisher Scientific; 52-821-0), the cells were fixed in a 1:1 mixture of methanol and acetone at −20°C for 10 minutes. Samples were washed three times with PBS containing 0.05% Tween 20, blocked with TBS containing 1%BSA (OmniPur BSA; EMD Millipore) for 1 hour at room temperature, and then incubated overnight at 4°C with primary antibodies diluted in the blocking buffer. The primary antibodies included acetylated α-Tubulin FITC (Santa Cruz Biotechnology; sc-23950 FITC), in-house anti-KIF7 polyclonal rabbit antibody, and in-house anti-GLI2 guinea pig antibody. KIF7 and GLI2 antibodies were labeled with Dylight^TM^650 and 488, respectively, using the DyLight™ Microscale Antibody Labeling Kits (Thermo Fisher#84536). Samples were mounted with ProLong Diamond Antifade Mountant (Thermo Fisher; P36961) for imaging.

### Sample preparation for TIRFM-based live cell imaging

NIH3T3 cells were seeded onto a 35mm No. 1.5 uncoated glass-bottom coverslip (Mattek; P35G-1.5-1.0-C). After 24 hours of seeding, when cells reached ∼ 80% confluency, the medium was replaced with pre-warmed low-serum DMEM for an additional 24 hours of incubation to induce ciliogenesis. On the imaging day, Halo-GLI2 was labeled using Janelia FluorX 549 HaloTag® Ligand (gift from Lavis Lab at HHMI, Janelia campus) at a final concentration of 200 nM for 1 hour. For imaging axoneme, 1 µM SiR-Tubulin was added for 1 hour prior to imaging. Cells were washed three times with pre-warmed DPBS, replenished with 1 ml of pre-warmed low-serum DMEM, and transferred to the TIRFM live-imaging chamber for imaging. To prevent photobleaching, antioxidant reagents, pyranose oxidase from Coriolus spp (Sigma; P4234; 100x: 70 mg/ml) and catalase from bovine liver (Sigma; C1345; 100x:3.6 mg/ml), were used. SAG21k (TOCRIS BIOSCIENCE), a high-affinity derivative of the SMO agonist, was added after the start of imaging to a final concentration of 1 uM for pathway activation.

### TIRF microscopy

#### Instrument setup

For live-cell imaging, a Nikon Ti2 inverted microscope equipped with a 100x/1.49 oil-immersion TIRFM objective lens was used. A 561 nm laser was employed for Halo-GLI2-JFX549 imaging, a 647 nm for SiR-Tubulin, and a 488 nm for mNeonGreen-IFT88 or SSTR3. The evanescent field penetration depth was set to 100-200 nm. The live-imaging chamber was maintained at 37°C and 5% CO_2,_ with a Zyla sCMOS camera used for multichannel imaging.

#### Selection of the primary cilium and acquisition parameters

Cilia were selected for live imaging based on three key criteria: (1) positivity for SSTR3-mNeonGreen or SiR-Tubulin labeling, (2) localization beneath the cell body, parallel to the coverslip with limited mobility during the entire imaging period, and (3) the presence of a faint signal of Halo-GLI2-JFX549 at the cilium tip before Hh signaling pathway activation. For long-duration TIRFM imaging, cells were imaged for 5-15 minutes before adding SAG21k, followed by continuous imaging for over 2 hours with 2-10-minute intervals between frames. For fast-acquisition imaging, images were acquired every 200-700 ms for up to 3 minutes per cilium. Cilia locations were pre-marked based on the cilia selection criteria above. After the addition of SAG21k, only cilia with observable GLI2 signal along the ciliary shaft - indicating that they were likely in the concentration phase-were selected for imaging

### Inhibitable motor assay

NIH3T3 Kif3a-/- kif3b-/- cells expressing i3A3B were seeded onto coverslips. After 24 hours, the culture medium was replaced with a starvation medium (DMEM containing 1% FBS). Following 48 hours of starvation, cells were treated either with vehicle (0.1% ethanol) or with 1 μM SAG21k, in combination with either DMSO or 50 nM B/B homodimerizer (Clontech). After 30 minutes of treatment, cells were fixed and stained with anti-IFT88 rabbit ab, anti-GLI2 guinea pig ab, and anti-acetylated α-tubulin FITC antibody. During analysis, cilia exhibiting irregular or “stubby” architecture were excluded using the “Analyze Particle” and “Thresholding” tools in FIJI.

### Image processing and data analysis

#### Kymographs

Movies were analyzed using FIJI. The KymoAnalyzer v1.01 plugin^70^ was used to generate kymographs and subsequently analyze the number, direction, and velocity of moving particles. To better visualize the trafficking along the ciliary shaft, kymographs were derived from line ROIs drawn along shaft, deliberately excluding oversaturated tip signals.

#### Intensity analysis

Custom codes in FIJI and MATLAB were employed to integrate the tip and whole cilia intensity of GLI2. Our MATLAB script automatically identified the start and end time points of the concentration phase.

#### Statistical analysis

GraphPad Prism was used for graph generation and statistical analysis.

### Star Table

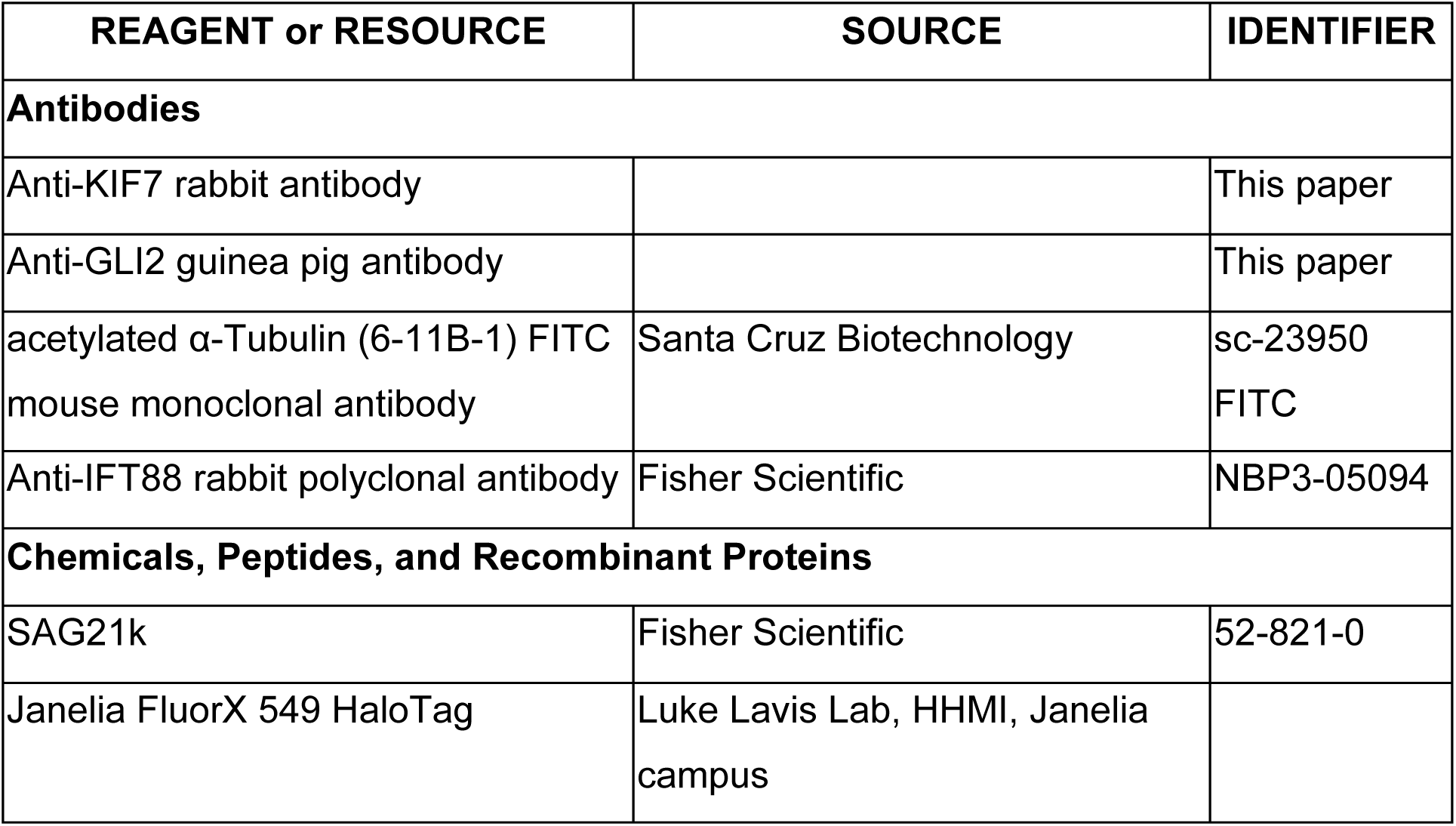
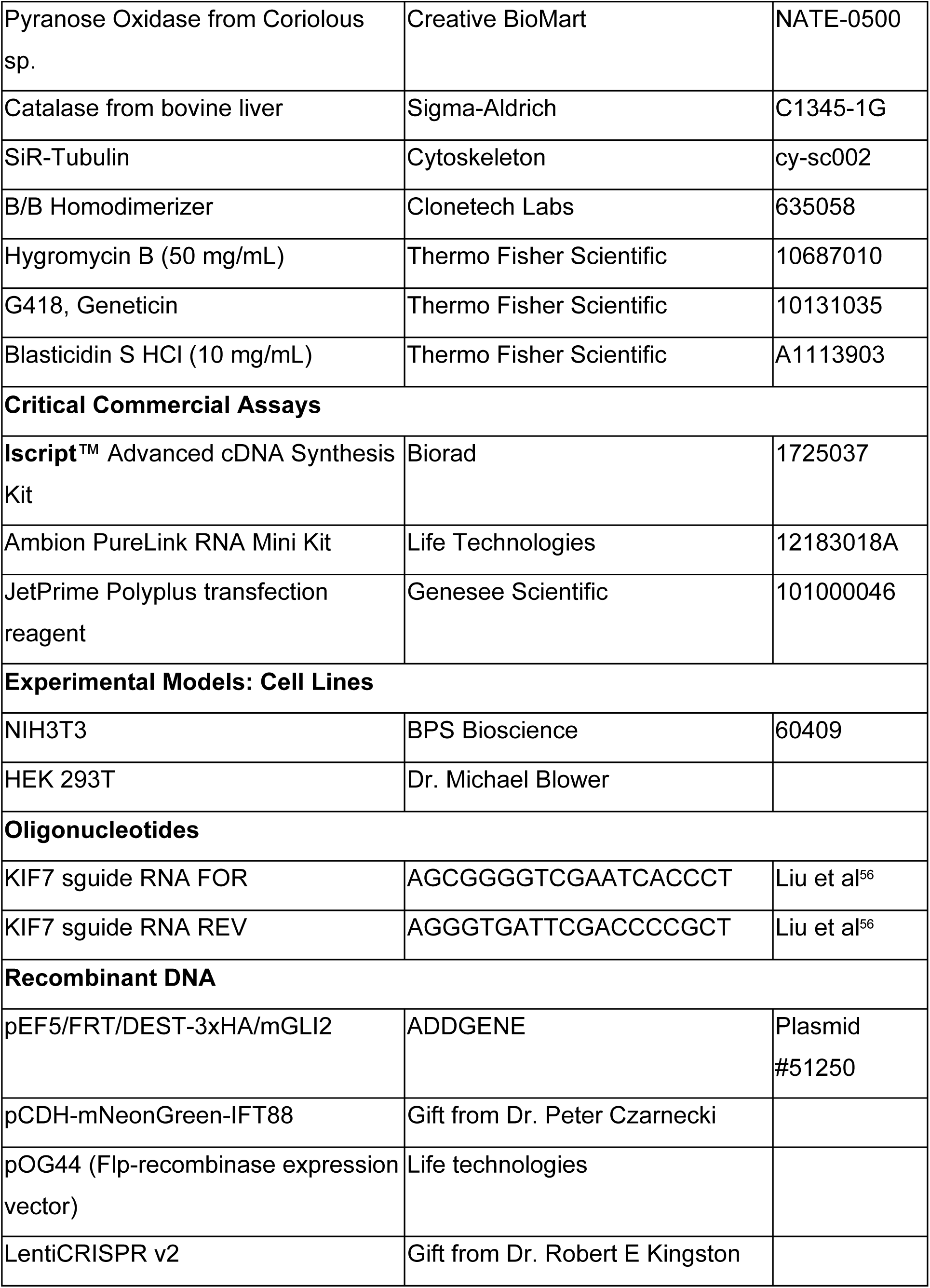
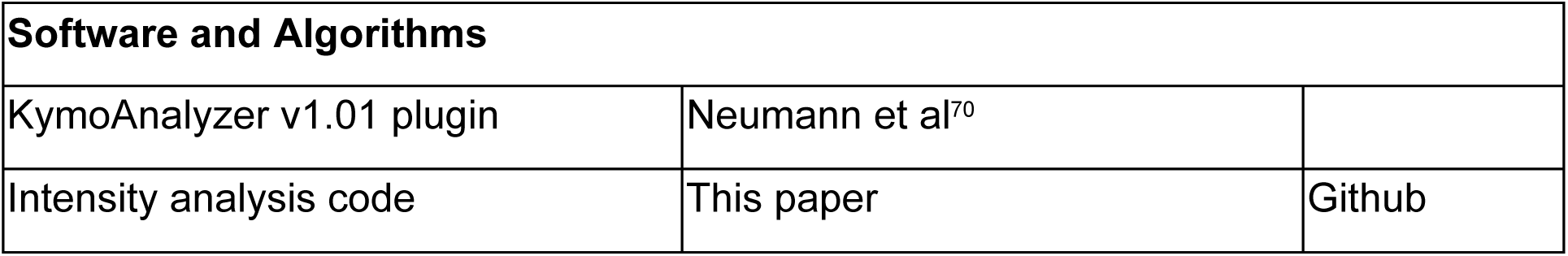

